# PIMO: Pathway-based Interpretable Multi-Omics interactions for multi-omics integration

**DOI:** 10.64898/2026.01.27.702136

**Authors:** Sai Phani Parsa, Sai Chandra Kosaraju, Euiseong Ko, Beomsu Baek, Tesfaye B. Marsha, Mingon Kang

## Abstract

**Motivation:** Modeling inter-omics interactions across multiple molecular levels is critical for deciphering the mechanisms underlying complex diseases. Epigenomic and structural alterations, such as DNA methylation and copy number alterations, modulate gene expression and collectively influence disease progression and patient survival outcomes. Despite advancements in deep learning-based multi-omics analysis, gene-level interactions of inter-omics have been seldom considered, due to combinational complexity and power, which limits interpretability and mechanistic insight.

**Results:** We propose a Pathway-based Interpretable deep learning Multi-Omics interaction model, PIMO, that explicitly captures regulatory effects across omics layers. Experiments on multiple TCGA cancer datasets showed that PIMO consistently outperformed state-of-the-art baselines in survival analysis, up to 13% increase in the C-index. PIMO provides biologically interpretable analyses that identify important pathways, genes, and inter-omics interactions with DNA methylation and copy number alterations.

**Availability and implementation:** The source code and data is available at https://github.com/datax-lab/PIMO.

## 1. Introduction

Characterization of inter-omics interactions of multilayered biological systems is essential for understanding complex disease pathogenesis that arises from cascades of molecular events across multiple omics layers, driven by interactions among transcriptomic, epigenomic, proteomic, and metabolomic processes (Kang et al., 2022; Subramanian et al., 2020). The inter-omics interactions of the multiple molecular layers correspond to regulatory processes that determine cellular function and influence patient outcomes (Chaudhary et al., 2018; Sharma et al., 2024; Liu et al., 2024). For instance, transcriptomic regulation occurs within structured biological systems, where regulatory signals propagate through coordinated pathways and are influenced by upstream epigenomic and genomic alterations. DNA methylation regulates transcriptional activity by modulating chromatin accessibility and gene regulatory programs. Regulatory changes in mutations that affect DNA methylation have been linked to cancer and other complex disorders (Farsetti et al., 2023; Zhang et al., 2020). Copy number alterations mask transcriptional outputs by changing gene dosage through regional amplifications and deletions, which results in systematic changes in transcript abundance (Louhimo and Hautaniemi, 2011; Pinkel and Albertson, 2005).

Deep learning–based multi-omics integration models have made significant progress in translating feature representations from multi-omics datasets, enabling comprehensive biological insights into molecular processes for precision medicine (Kang et al., 2022; Ballard et al., 2024). Multi-omics integration approaches can be categorized into two categories: (1) combining single-omics predictions obtained from separately trained models for each modality, and (2) training a unified network that learns a concatenated representation shared across multiple modalities (Li et al., 2022; Sucre et al., 2025; Nikolaou et al., 2025; Qu et al., 2025b). First, integrating prediction scores from different omics models effectively merges omics-specific signals for downstream tasks. Individual multilayer perceptron models have been trained on each individual omics dataset for survival analysis, and their outputs have then been merged to generate integrated survival predictions (Sucre et al., 2025). A weighted ensembling strategy has been proposed to aggregate prediction scores generated by single omics models, allowing different contributions from each modality (Nikolaou et al., 2025). Such models are challenging to interpret and are vulnerable to biases introduced by individual omics modalities.

Alternatively, latent representation–level integration offers greater interpretability than prediction-level aggregation models. These models often implement inter-omics latent representation integration within a unified network, for multi-omics integration (Farsetti et al., 2023; Uyar et al., 2025). For example, PCLSurv introduced a concatenation framework built on an encoder–decoder architecture to generate modality-specific embeddings for predicting patient risk scores (Li et al., 2025). DeepKEGG used self-attention mechanisms to derive latent features from each individual omics modality and then combined these representations through concatenation (Lan et al., 2024). MOGONET adopted a graph-based information-sharing strategy, in which separate graph convolutional networks were trained for each omics modality, and inter-omics correlations were modeled for cancer classification. (Wang et al., 2021). GraphPATH built a biologically informed graph-based network from DNA methylation and copy number alteration, and then applied multi-head attention for tumor classification(Ma and Wang, 2024).

Despite advances in multi-omics integration, interactions between different omics layers have been seldom considered. The inter-omics interaction analyses include computational complexity, small sample sizes, and the nonlinear nature of the interactions. Generating pairwise interactions between omic layers in such high-dimensional settings tends to cause overfitting. In addition, the complex structure of each data modality can yield a large number of biologically irrelevant feature pairs, which may ultimately degrade both the predictive performance of the model and its interpretability.

Pathway-based modeling improves interpretation and constrains high-dimensional molecular analysis by focusing on biologically predefined gene groups. As it narrows the potential search space for molecular interactions, this approach helps to prevent overfitting and strengthens statistical reliability, all while retaining connections that matter for patient outcomes (Hao et al., 2018). Interpretations derived from pathway-structured representations facilitate biologically meaningful understanding of survival analysis and classification (Oh et al., 2021; Elmarakeby et al., 2021).

In this study, we introduce PIMO, a Pathway-based, interpretable Multi-Omics interactions model designed to capture relationships among transcriptomics, DNA methylation, and copy number alterations. PIMO employs an interaction-aware network inspired by cross-attention mechanisms, leveraging key and query representations. More precisely, PIMO represents inter-omics interactions by using a key representation for transcriptomic features together with query representations for DNA methylation and copy number alterations. To ensure computational tractability and biologically interpretable results, PIMO restricts inter-omics interaction pairs to occur within the same pathway. We assessed the performance of PIMO for survival analysis under several experimental scenarios: (1) repeated Monte Carlo cross-validation with ten independent runs on multiple TCGA cancer datasets, (2) external validation using an independent METABRIC-BRCA dataset, and (3) investigation of gene-level inter-omics interaction contributions to survival outcomes.

## 2. Methods

### 2.1. Model Overview

PIMO integrates gene expression, DNA methylation, and copy number alterations (CNA) data to model inter-omics relationships at the gene level. For each gene, expression features are encoded as key representations, while methylation and CNA features act as queries. A cross-attention–inspired mechanism learns regulatory patterns from these representations. Gene features are then grouped within each biological pathway and summarized to form pathway representations. These representations combine original gene signals with learned interaction patterns, enabling interpretation at the pathway, gene, and interaction levels. The final pathway representations support downstream survival analysis.

### 2.2. Model Architecture

The PIMO architecture is composed of four components: (1) a pathway-based multi-omics interaction layer, (2) a pathway representation layer, and (3) convolutional layer, and (4) output layers, as shown in Fig. 1. First, in the pathway-based multi-omics interaction layer, gene-level interactions between DNA methylation (*d*) and gene expression (*g*), as well as between copy number alterations (*c*) and gene expression, are first computed as

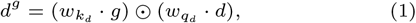

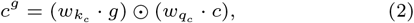

where 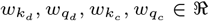 are learnable, gene-specific scalar weights that capture how gene expression interacts with the associated epigenomic and structural alterations. The resulting *d*^*g*^ and *c*^*g*^ respectively denote the gene-level interactions of DNA methylation and CNA on gene expression. Based on these interactions, a gene-level representation (***ϕ***) for the *i*^*th*^ gene is constructed by concatenating its gene expression value with its interaction features as

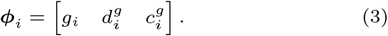

where ***ϕ***_***i***_ denotes the gene-level interaction features for the *i*^*th*^ gene, containing gene expression *g*_*i*_, interactions between DNA methylation and gene expression 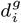 and interactions between CNA and gene expression 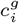 as shown in the Fig. 1B.

**Fig. 1.**
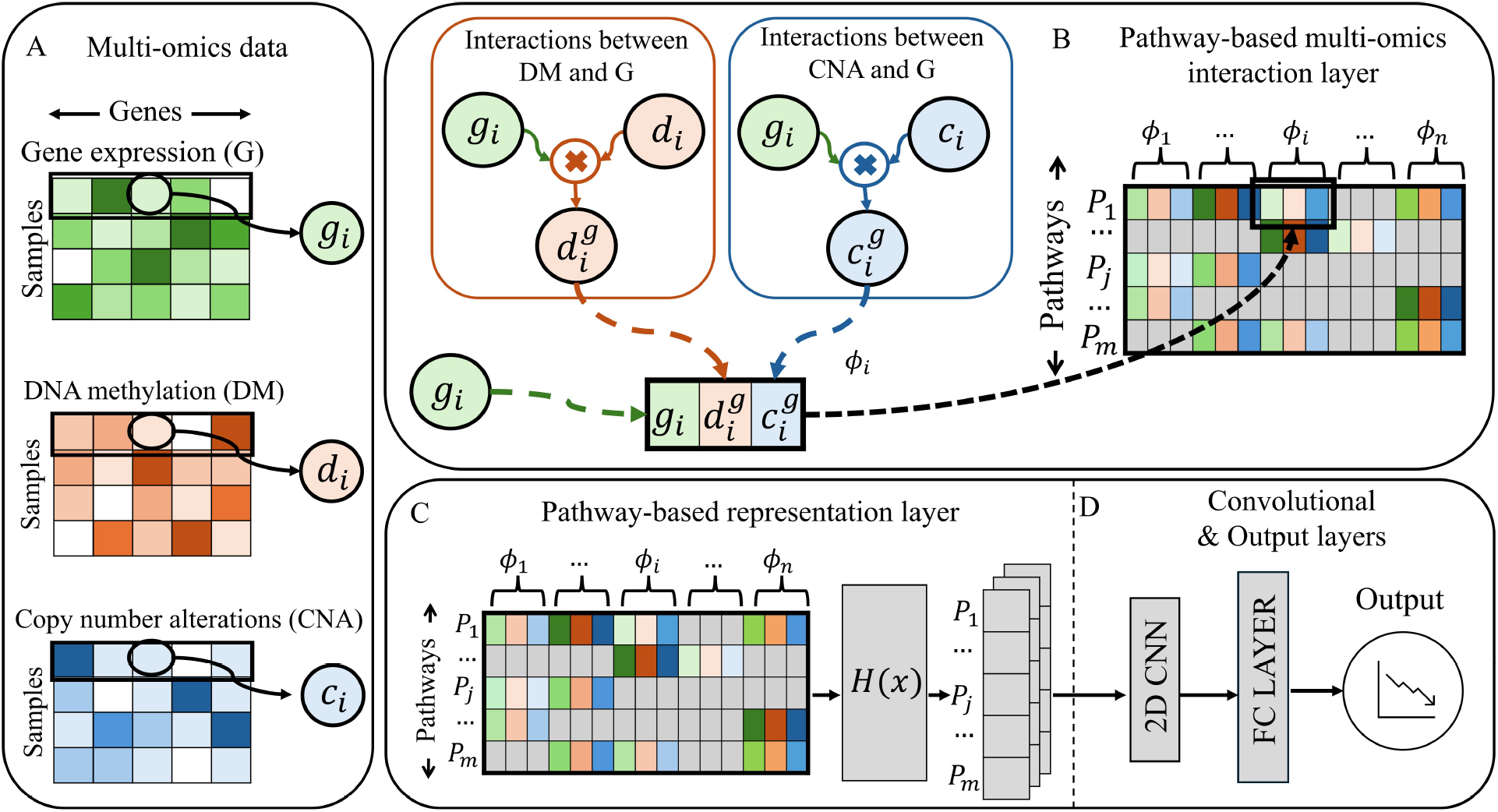
Overview of the proposed PIMO architecture. (A) **Multi-omics inputs:** For each gene *i*, gene expression (*g*_*i*_), DNA methylation (*d*_*i*_), and copy number alteration (*c*_*i*_) values are provided as gene-level inputs. (B) **Pathway-based multi-omics interaction layer:** Within each pathway, gene expression features are combined with DNA methylation and CNA through learnable gene-wise interaction functions, producing interaction features 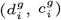 that are concatenated with *g*_*i*_ to form unified gene representations ***ϕ*** . (C) **Pathway-based representation layer:** Gene-level representations are aggregated within pathways and transformed using a learnable function *H*(*x*) to obtain pathway-level representations. (D) **Convolutional and output layers:** Pathway representations are further modeled using a 2D convolutional layer followed by fully connected layers to produce the final output.

Pathway structure for the *j*^*th*^ (1 ≤ *j* ≤ *m*) pathway is established through a binary pathway membership mask ℳ ^(*j*)^ ∈{0, 1}^*n*^, where *m, n* denotes the total number of pathways and genes respectively. 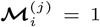 if the *i*^*th*^ gene belongs to the *j*^*th*^ pathway, and 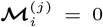otherwise. Using this pathway membership mask, gene-level representations for the *j*^*th*^ pathway are aggregated by masking and horizontally stacking as

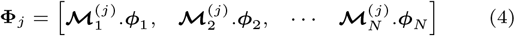

where **Φ**_***j***_ ∈ ℜ^1×3*n*^, since each gene-level representations (***ϕ***_***i***_) is comprised of three elements.

For a sample *k*, consisting of *m* pathways and *n* genes, gene-level representations at the sample-level are obtained as:

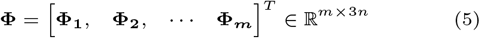

Pathway ordering in **Φ** is determined by representing each pathway as a fixed-length feature vector obtained from gene-level multi-omics data aggregated at the pathway level, followed by computing pairwise Pearson correlation coefficients between pathways across all samples. Pathways are then sequentially arranged by placing highly correlated pathways in close proximity, resulting in a fixed, correlation-aware ordering that is shared across all samples. This ordering preserves similarity relationships among pathways while providing a consistent structural organization of pathway representations for subsequent modeling stages (Oh et al., 2021).

Second, the pathway representation layer (Fig. 1C) computes pathway representations (**P**) by convoluting *H*(*x*) with **Φ** as

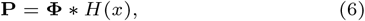

where *H*(*x*) denotes *γ* number of learnable linear filters, where each filter is of dimension *n* × 1.

Third, the pathway-pathway interactions (Fig. 1D) are obtained by applying a 2D convolutional layer on pathway representations (**P** ∈ ℜ^1×*m*×*γ*^ ). Finally, the resulting features are passed through fully connected layers to learn higher-level abstract representations for downstream tasks. For survival analysis, PIMO produces a prognostic index (PI) for each patient, which serves as the risk score in a Cox proportional hazards (Cox-PH) regression model, with model parameters optimized by minimizing the negative partial log-likelihood. In addition to survival analysis, the framework can be readily extended to classification tasks by modifying the final output layer to a softmax or sigmoid activation, enabling the prediction of discrete clinical outcomes.

### 2.3. Model interpretation

PIMO provides model interpretation at three stages: pathway-level (*I*^*p*^), gene-level (*I*^*g*^), and inter-omics interaction-level (*I*^*m*^). The pathway importance scores (*I*^*p*^) capture the relative contribution of biological pathways to model predictions, enabling the identification of key signaling processes associated with patient outcomes. Gene importance scores (*I*^*g*^) measure the influence of individual genes within each omics layer, highlighting potential molecular drivers and candidate biomarkers. Inter-omics interaction importance scores (*I*^*m*^) quantify inter-omics regulatory effects, revealing how epigenomic and systematic alterations modulate transcriptomic activity. These three measures together provide a comprehensive, interpretable view of disease-associated molecular mechanisms.

#### 2.3.1. Pathway importance scores

The importance of a pathway *P*_*j*_ is quantified by assessing its contribution to the model prediction across latent feature maps derived from the pathway representation layer. Specifically, for each pathway *j* and latent feature map *l*, we compute the gradient of the model output with respect to the corresponding activation map 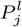 as

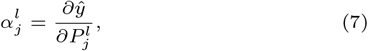

where *ŷ* denotes the prognostic index and 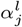 captures how strongly the *l*^*th*^ latent representation of pathway *P*_*j*_ influences the prediction. These gradients capture the contribution of each pathway-specific feature map to the model output.

The pathway-level importance score is then obtained by aggregating contributions across all *γ* latent feature maps as

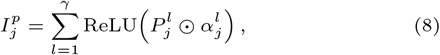

where the element-wise product weights each pathway activation by its corresponding gradient contribution, and the ReLU operation retains only positively contributing features. This formulation emphasizes pathways that consistently increase predicted risk across latent spaces.

A binomial statistical test is performed to evaluate the statistical significance of pathway importance scores. For each sample, the pathway importance is treated as a Bernoulli trial that records whether the pathway attains a positive importance score. A binary indicator for sample *k* and pathway *j*, are defined as

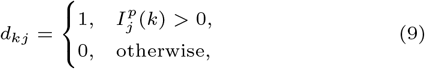

where 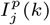 denotes the importance score for pathway *j* in sample *k*. A global baseline positive attribution rate is calculated over all pathways and samples, reflecting the expected likelihood of encountering a positive attribution for any given pathway. The global baseline positive attribution rate is computed as

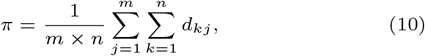

representing the expected probability of observing a positive attribution across all pathways and samples. The pathway-specific positive rate can be estimated as

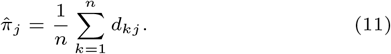

In the binomial test, the null hypothesis is formulated as *H*_0_ : *π*_*j*_ ≤ *π*, which evaluates whether the attribution frequency of pathway *j* is greater than the global baseline positive attribution. Pathways for which *π*_*j*_ *> π* are deemed statistically significant. Finally, all *p*-values are corrected for multiple testing using the Benjamini-Hochberg FDR procedure with a threshold of 0.05.

#### 2.3.2 Gene and Interaction Importance Scores

Gene importance is computed by propagating pathway-level importance scores to individual gene expression features as

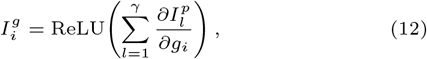

where *g*_*i*_ denotes the expression value of the *i*^*th*^ gene. Similarly, interactions importance DNA methylation 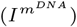 and CNA 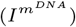 are computed as:

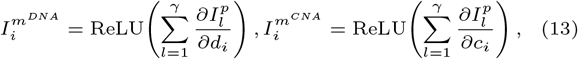

respectively.

Statistical significance of *I*^*g*^ and *I*^*m*^ is assessed using a binomial test following the same procedure as the pathway-level analysis. The resulting significant genes are subsequently mapped to their corresponding pathways to identify pathway-specific molecular drivers.

## 3. Results

In this section, we present the empirical assessment of PIMO across multiple experimental settings, including (1) survival analysis performance evaluated using ten independent Monte Carlo cross-validation (MCCV) trials on multiple TCGA cancer datasets, (2) model generalizability evaluated through external validation, and (3) ablation studies assessing the contribution of pathway-based gene-wise inter-omics interaction modeling.

### 3.1. Data preprocessing

We evaluated the proposed model using multiple TCGA cancer datasets, including breast invasive carcinoma (BRCA), brain lower-grade glioma (LGG), lung adenocarcinoma (LUAD), liver hepatocellular carcinoma (LIHC), and kidney renal clear cell carcinoma (KIRC). These datasets were obtained from the cBioPortal for Cancer Genomics (cBioPortal for Cancer Genomics, 2012). Detailed information on sample sizes for each cohort is provided in Table 1.

**Table 1.**
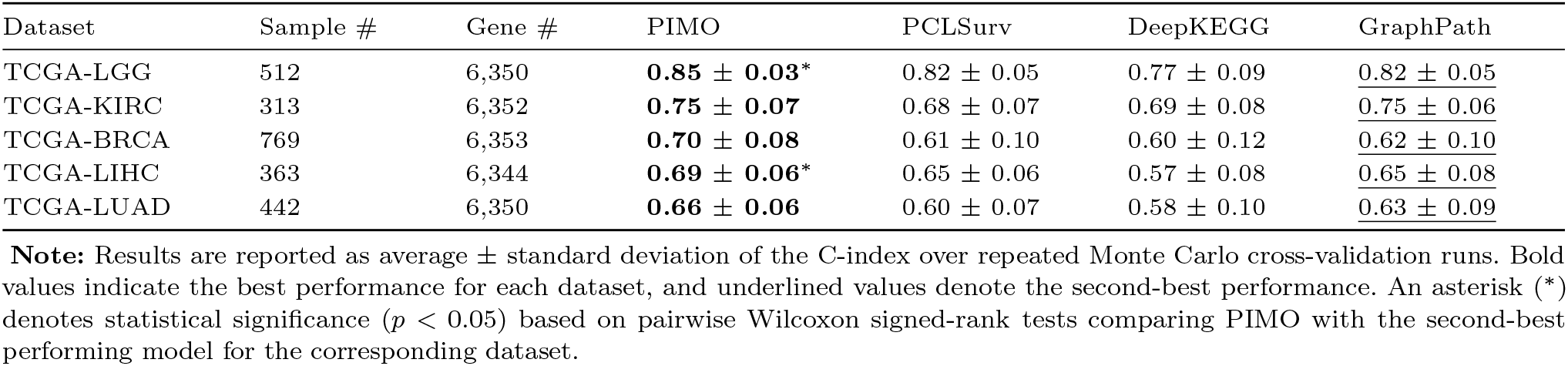
Survival analysis performance across TCGA datasets measured by C-index (average *±* std).

We incorporated biological pathway information from the Kyoto Encyclopedia of Genes and Genomes (KEGG) pathway database (Kyoto Encyclopedia of Genes and Genomes (KEGG), 2025), which compiles molecular interaction and reaction networks. We first retrieved 365 human (*Homo sapiens*) pathways from KEGG. To reduce bias, we excluded human disease–specific KEGG pathways, leaving 257 pathways. To derive robust and biologically meaningful pathway-level representations, we additionally filtered out pathways containing fewer than 15 genes or more than 300 genes, resulting in a final set of 231 pathways.

For each dataset, we defined the reference gene set as the genes available in the gene expression modality. To maintain consistency of genes across modalities, we used this same set of genes as the reference for all modalities. We incorporated DNA methylation and CNA data only for these reference genes, imputing a value of 0 whenever DNA methylation or CNA measurements were missing for a reference gene. Finally, we standardized all gene-level features within each modality to have zero mean and unit variance before training the model.

### 3.2. PIMO’s survival analysis performance on TCGA cancer datasets

We compared PIMO against three state-of-the-art (SOTA) multi-omics models, including DeepKEGG (Lan et al., 2024), PCLSurv (Li et al., 2025), and GraphPath (Ma and Wang, 2024). These benchmark models represent distinct multi-omics integration strategies, ranging from pathway-informed representations to feature- and graph-based fusion mechanisms. Although DeepKEGG and GraphPath were originally evaluated on tasks such as recurrence prediction and cancer status classification, we adapted them for survival analysis by replacing their respective final layers with a Cox layer. We used Optuna (Akiba et al., 2019) to perform hyperparameter tuning for all models by minimizing the validation loss.

We performed ten independent Monte Carlo cross-validation (MCCV) experiments with an 80%/10%/10% split for training, validation, and test sets, respectively, to ensure the reproducibility of our findings. Survival analysis performance was evaluated using the Concordance Index (C-index). For PIMO, the optimal hyperparameter configuration uses 32 kernels in the pathway representation layer (*γ* = 32) and 4 kernels in the 2D convolutional layer. In each Monte Carlo cross-validation run, we tuned the learning rate within the range 1*e*∈^5^ to 1*e*^−4^ and used a multiplicative learning-rate scheduler that decreased the learning rate by a factor between 1*e*^−1^ and 9*e*^−1^ at regular intervals of 50, 100, 150, or 200 epochs, as defined during hyperparameter optimization. We trained PIMO with the Adam optimizer and applied early stopping based on the validation loss to mitigate overfitting, selecting the final model as the checkpoint with the best validation performance.

PIMO consistently achieved the higher average C-index across all evaluated TCGA cancer datasets, with GraphPath generally emerging as the strongest competing baseline. Across the TCGA-LGG and TCGA-LIHC datasets, PIMO significantly outperformed the SOTA methods, achieving average C-indices of 0.85 *±* 0.03 and 0.69 *±* 0.06, respectively, with *p*-value *<* 0.05 based on Wilcoxon signed-rank tests. These outcomes correspond to relative improvements of 3 ∼ 21% over GraphPath, DeepKEGG, and PCLSurv.

For both TCGA-BRCA and TCGA-LUAD, PIMO obtained the highest average C-index values of 0.70 *±* 0.08 and 0.66 *±* 0.06, respectively, surpassing the state-of-the-art approaches, whose second-best performances were 0.82 *±* 0.05 and 0.63 *±* 0.09. On TCGA-KIRC, which has the smallest sample size, PIMO achieved the joint-best performance of 0.75 *±* 0.07, matching GraphPath (0.75 *±* 0.06) and exceeding PCLSurv and DeepKEGG by at least 7%. Furthermore, Table 1 and Fig. 2 illustrate the performance of PIMO in comparison with state-of-the-art benchmark models.

**Fig. 2.**
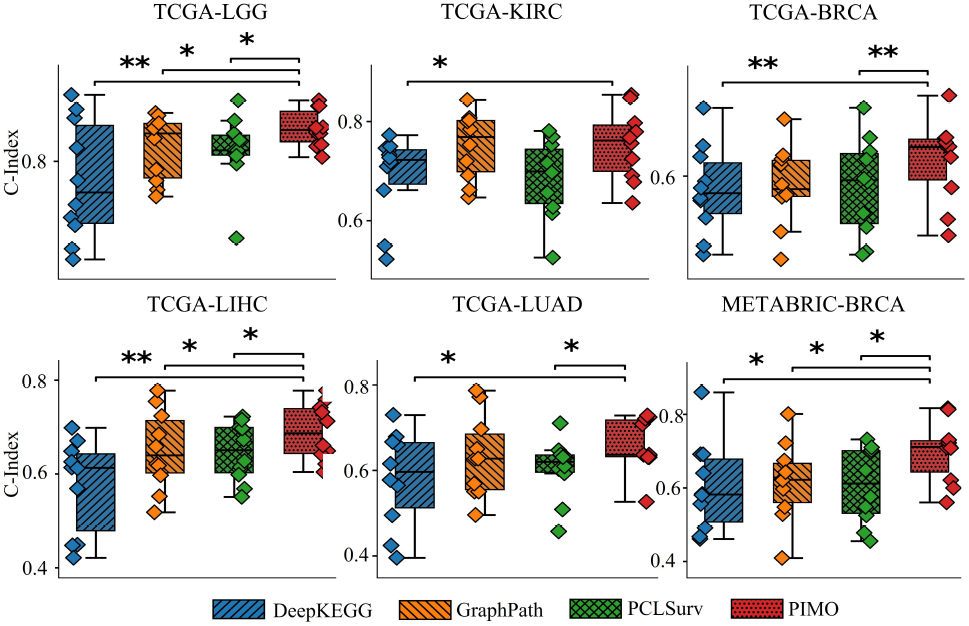
Box plots summarize the distributions of C-index values from ten repeated Monte Carlo cross-validation runs for survival analysis across multiple TCGA cancer datasets (KIRC, LIHC, LUAD, LGG, BRCA) and the METABRIC (BRCA) dataset. Individual points shown as overlaid strips represent C-index values from each cross-validation run. The central line in each box denotes the median C-index, while the box spans the interquartile range (25th-75th percentiles). Statistical significance is indicated using star notation based on pairwise Wilcoxon signed-rank tests, where ** denotes *p <* 0.01 and * denotes *p <* 0.05.

### 3.3. External validation using independent METABRIC-BRCA dataset

To further examine the generalizability of PIMO, we evaluated performance on an external dataset from the Breast Cancer International Consortium Molecular Taxonomy dataset (METABRIC-BRCA). This dataset consists of 1,416 breast cancer patients, including 830 uncensored and 586 censored samples, retrieved from cBioPortal for Cancer Genomics (2012). PIMO obtained the highest average C-index (0.58 *±* 0.01), outperforming GraphPath (0.56 *±* 0.02), PCLSurv (0.53 *±* 0.04), and DeepKEGG (0.56 *±* 0.03) (Table 2), demonstrating its superior generalization capability. In addition, PIMO exhibited the smallest standard deviation among all methods, highlighting its reliability and statistical robustness.

**Table 2.**
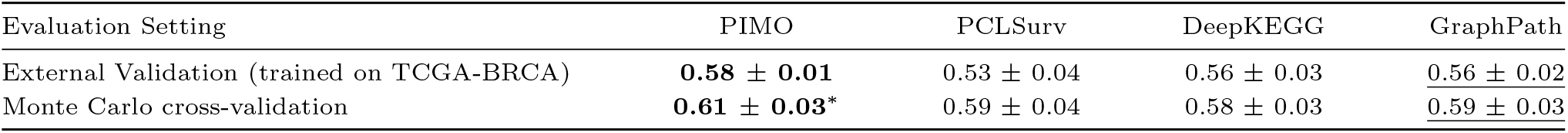
Survival analysis performance on the METABRIC-BRCA dataset under external validation and Monte Carlo cross-validation, measured by C-index (average *±* std).

We additionally assessed the model’s robustness and versatility by training PIMO and the other benchmark methods with MCCV on the relatively large METABRIC-BRCA dataset (1416 samples), using a training strategy analogous to that used for the TCGA datasets. PIMO consistently outperformed all benchmarks by at least 3%. PIMO obtained an average C-index of (0.61 *±* 0.03) with a *p*-value *<* 0.05. Table 2 presents the performance comparison on METABRIC-BRCA; collectively, these findings demonstrate PIMO’s strong generalizability and efficiency across multiple datasets.

### 3.4. Interaction effects on predictive performance

To investigate the impact of inter-omics interactions on model predictions, we modified the PIMO architecture by disabling the interaction components, replacing 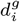 with *d*_*i*_ and 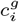 with *c*_*i*_ in *ϕ*_*i*_ in equation 3 (refer to 2.2), and then directly comparing this ablated variant to the original PIMO model. We conducted this evaluation on the TCGA-BRCA and TCGA-LGG datasets, and in both cases, the interaction-enabled PIMO outperformed the version without interactions. On TCGA-BRCA, PIMO with interactions achieved an average C-index of 0.70 *±* 0.08, corresponding to a 7% increase compared to PIMO without interactions. Table 3 presents the comparison between PIMO models with and without interactions. These findings suggest that gene-level inter-omics interactions can meaningfully enhance patient-outcome prediction.

**Table 3.** Interaction effects of pathway-based gene-wise inter-omics modeling on survival analysis performance, measured by C-index (average *±* std).

| Dataset | Gene-wise inter-omics interactions | C-index (average ± std) |
| --- | --- | --- |
| TCGA-BRCA | ✗ | 0.65 ± 0.10 |
|  | ✓ | **0.70 ± 0.08** |
| TCGA-LGG | ✗ | 0.83 ± 0.05 |
|  | ✓ | **0.85 ± 0.03** |

**Note:** A check mark (✓) indicates that gene-wise inter-omics interaction modeling is enabled, while a cross (✗) indicates that it is disabled.

### 3.5. PIMO interpretation findings

For model interpretation, we retrained PIMO on the entire TCGA-BRCA dataset using hyperparameters that were consistently identified as optimal across the ten experiments. We then calculated *I*^*p*^, *I*^*g*^, and *I*^*m*^ on this final model to assess pathway, gene, and interaction importance, respectively. Furthermore, the resulting PIMO interpretation outcomes are subsequently correlated with the original data to uncover relevant pathways and gene functions associated with disease prognosis.

#### 3.5.1. Pathway importance identified by PIMO

PIMO identified 11 out of 231 pathways as important and statistically significant for survival outcomes (as described in 2.3). Table 4 presents these 11 pathways together with supporting biological evidence for reference. Nearly half of the important pathways detected by PIMO have been previously reported in the literature as being relevant to disease prognosis. Even after excluding directly disease-specific pathways (as in 3.1) to reduce bias, PIMO’s detection of biologically supported pathways demonstrates its potential to identify disease-specific pathways.

**Table 4.** Top 11 PIMO-identified pathways associated with cancer survival, ranked by pathway importance scores, with corresponding *p*-values and supporting references.

| Pathway | Imp. Score | $p$-value | PMID |
| --- | --- | --- | --- |
| Antigen processing and presentation | $9.8e^{-3}$ | $2.09e^{-4}$ | 32070368 |
| TGF-β signaling | $5.2e^{-3}$ | $2.62e^{-2}$ | 23022998 |
| RLR signaling | $3.8e^{-3}$ | $7.70e^{-5}$ | |
| Cytosolic DNA-sensing | $3.2e^{-3}$ | $2.09e^{-4}$ | |
| Ovarian steroidogenesis | $2.7e^{-3}$ | $1.53e^{-2}$ | |
| Vasopressin-regulated water reabsorption | $2.0e^{-3}$ | $2.09e^{-4}$ | 33968066 |
| Folate transport and metabolism | $1.0e^{-3}$ | $3.00e^{-6}$ | |
| Virion–Lassa virus and SFTS virus | $9.4e^{-4}$ | $2.95e^{-3}$ | |
| Phenylalanine metabolism | $7.6e^{-4}$ | $1.77e^{-2}$ | 30830383 |
| Butanoate metabolism | $4.7e^{-4}$ | $2.09e^{-4}$ | 37495653 |
| Selenocompound metabolism | $4.2e^{-4}$ | $1.53e^{-2}$ | |

Antigen Processing and Presentation (APP) identified as the most important pathway by PIMO, exhibiting an importance score of *I*^*p*^ = 9.8*e*^−3^ and a p-value of 2.09*e*^−4^. APP has been associated with breast cancer due to its relationship with glycolysis activity. Elevated glycolysis enhances the activation of immune-related pathways in which APP is involved, potentially influencing cancer outcomes (Table 4).

The TGF-*β* signaling pathway is the second-most-important pathway identified by PIMO. It is linked to cancer development by facilitating tumor cell invasion and migration. TGF-*β* has also been recognized as a driver of tumorigenesis, promoting tumor progression (Table 4). The Vasopressin-regulated water reabsorption pathway identified by PIMO is among those that control CD2 expression, which, in turn, influences survival outcomes in breast cancer patients (Table 4). Interestingly, PIMO highlights Phenylalanine and Butanoate metabolism pathways in specific demographic patient subgroups, both of which are biologically linked to cancer progression. The phenylalanine metabolism pathway has been associated with cancer progression in a sub-group of Asian populations (Table 4), while butanone metabolism has been identified as a key cancer-related pathway in white populations (Table 4). Together, these findings suggest that PIMO can capture biologically meaningful patterns relevant to cancer prognosis.

To evaluate the prognostic relevance of the main PIMO-derived pathways, we divided the samples into two groups according to the median importance score for each pathway and compared survival outcomes using Kaplan–Meier analyses. For the grouping, we excluded samples with censored observations when the survival time was ≤ 60 months. We then used the log-rank test to determine whether survival differed significantly between the two groups and accounted for multiple comparisons by applying an FDR correction at 0.05 to the resulting *p*-values.

Nine out of the eleven important pathways had *p*-values *<* 0.05, indicating that the pathways identified as important by PIMO are robust predictors in the survival analysis. For example, the APP pathway showed a clear separation between the two groups, with a *p*-value of 2.09*e*^−4^, indicating that patients with higher APP pathway activation experience poorer survival, underscoring its clinical relevance (Fig. 3B). Fig. 3B shows the Kaplan–Meier plots for the important pathways identified by PIMO.

**Fig. 3.**
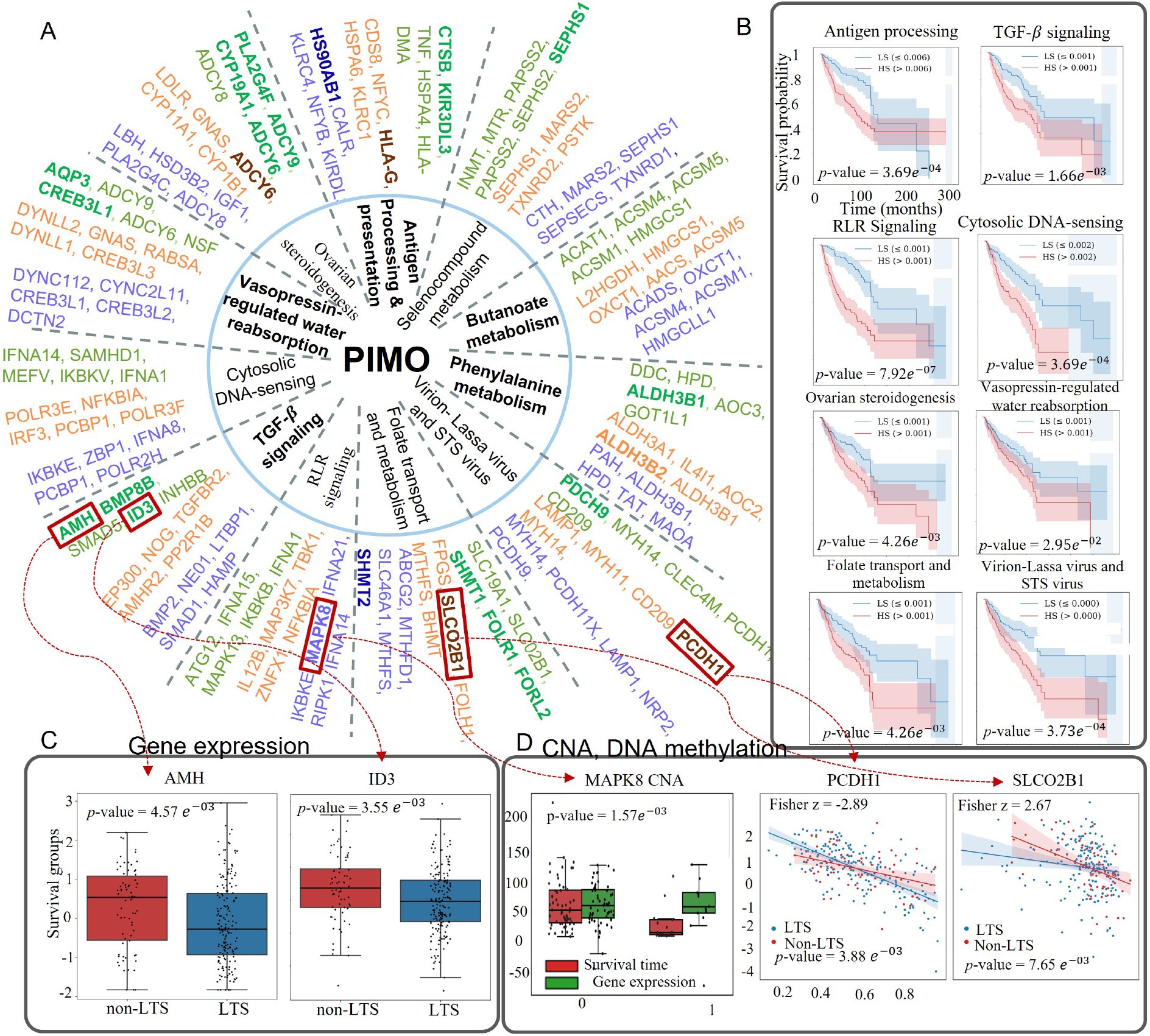
PIMO interaction findings of important pathways, genes, and interactions. (A) Overall, PIMO identified 11 pathways as significantly associated with disease prognosis. The five highest-ranking genes were chosen from these significant pathways. Green, orange, and blue labels denote the top five gene expression features, DNA methylation interactions, and CNA interactions, respectively. Genes and interactions shown in bold are those supported by existing biological literature. (B) Kaplan–Meier plots demonstrating the prognostic relevance of the important pathways. (C) Box plots of important gene-expression having differential distributions between LTS and non-LTS groups. (D) Differential distributions of important interactions of CNA and DNA methyalation.

#### 3.5.2. Gene-expression importance identified by PIMO

To assess gene importance (*I*^*g*^), we selected the top five most important genes for each of the 11 significant pathways. Over 15% of the key genes identified by PIMO are supported by biological evidence reported in the literature. PIMO identifies CTSB and KIR3DL3 as significant genes involved in the APP pathway. CTSB is implicated in regulating the APP pathway, which has been linked to poorer survival outcomes (Zhou et al., 2020). Overexpression of KIR3DL3 has been linked to more aggressive tumor progression (Montoyo-Pujol et al., 2025). Within the TGF-*β* pathway, PIMO identifies AMH, BMP8B, and ID3 as key genes associated with survival and high-risk prognostic markers (Wang et al., 2025; Katsuta et al., 2019; Qu et al., 2025a). ATG12 has been identified as the most significant gene within the RLR signaling pathway. Elevated ATG12 expression is associated with worse survival outcomes in BRCA (Zhang et al., 2023). SAMHD1 expression, which is associated with disease-free progression for HER2-enriched subtypes, is identified as an important gene in the Cytosolic DNA-sensing pathway (Kouvaraki et al., 2025). Additional genes, such as PLA2G4F, ACAT1, CYP19A1, ADCY6, CREB3L1, SHMT1, FOLR1, PCDH9, and SEPHS2, have been recognized as key genes within 11 significant pathways linked to tumor progression and patient survival (Tang et al., 2025; Kuldeep et al., 2023; Long et al., 2006; Wang et al., 2019; Wang and Jiang, 2024; Kim et al., 2014; Liu et al., 2020; Li et al., 2024; Cao et al., 2025). Fig. 3A shows the full list of genes identified by PIMO.

To investigate the distributional characteristics of PIMO-identified genes, we compared gene expression levels between long-term survivors (LTS) and non-LTS patients using the Mann–Whitney U test. Patients with survival times longer than 60 months were designated as LTS, whereas all remaining uncensored patients were categorized as non-LTS. Four genes (AMH, ID3, MAPK13, and ACAT1) showed clearly different distribution patterns between the two groups. Among these, AMH and ID3 are illustrated as box plots in Fig. 3C. These box plots reveal differences in the distributions of gene expression levels between the LTS and non-LTS groups, indicating shifts in both central tendency and variability across the two survival categories.

#### 3.5.3. Interaction importance of DNA-methyalation and Copy Number Alteration

Similar to gene expression, we determined the top-5 most important interactions (*I*^*m*^) for 11 significant pathways. PIMO identifies at least five interactions supported by biological evidence. Although the number of validated interactions is currently limited, we believe that future diagnostic efforts will increasingly focus on such interactions, prompting more biological studies in this area. In this context, PIMO’s findings can offer valuable insights to support these future investigations.

PIMO identified copy number alterations of HSP90AB1 as the most significant interaction within the AAP pathway. Copy number alterations in HSP90AB1 have been associated with poorer outcomes in breast cancer (Cheng et al., 2012). Similarly, overexpression of SHMT2 driven by copy number alterations was identified as a key interaction in the folate transport and metabolism pathway. Copy number alterations in SHMT2 have been linked to unfavorable prognosis and reduced relapse-free survival (Usman et al., 2023).

According to PIMO, HLA-G and ADCY6 are significant DNA-methylation and gene expression interactions for AAP and the ovarian steroidogenesis pathway, respectively. The regulatory effects of HLA-G and ADCY6 are linked to patient prognosis, particularly in the context of breast cancer progression and outcome. For example, HLA-G has been reported to be hypermethylated in inflammatory breast cancer tissues (Calanca et al., 2024), and hypermethylation-associated downregulation of ADCY6 is associated with improved prognosis (Li et al., 2020). PIMO also identifies ALDH3B2 as a significant DNA methylation–gene expression interaction within the phenylalanine metabolism pathway. Regulatory effects of ALDH3B2 DNA methylation have been implicated in membrane lipid biosynthesis in BRCA (Xu et al., 2022).

We further examined the underlying data distributions, analogous to those of gene expression, to explore patterns associated with copy number alteration (CNA). The CNA status of MAPK8 exhibits a notably distinct distribution. In Fig. 3D, we present the distributions of survival time and gene expression for samples with CNA values of 0 and 1; the accompanying *p*-value reflects an unadjusted comparison between these two CNA categories. This visualization underscores clear differences in both the central tendency and the variability of survival times across the CNA groups.

We additionally investigated DNA methylation–gene expression relationships in a way similar to the CNA-gene expression analysis. For two DNA methylation sites, PCDH1 and SLCO2B1, the scatter plots in Fig. 3D show differences in distribution between the two survival groups. The Fisher’s z-statistics and the corresponding *p*-values are also reported in Fig. 3D.

## 4. Discussion

In this study, we introduce an interpretable, interaction-aware deep learning model, named PIMO. PIMO explicitly represents gene-level inter-omics interactions of multi-omics data in a biological pathway. We assessed PIMO’s performance on multiple TCGA datasets. PIMO consistently surpassed existing benchmark methods in the predictive performance, indicating robust generalization across a wide range of cancer types. PIMO discovered significant pathways and genes and their interactions with DNA methylation and copy number alternation.

Examination of inter-omics interaction scores uncovered regulatory effects of DNA methylation on transcriptional activity, along with structure variation arising from copy number alterations. In-depth exploratory data analysis further revealed that the ID3 and AMH genes exhibited distinct expression patterns between long-term survivors (LTS) and non-LTS patients, indicating potential prognostic significance. Additionally, copy number alteration in MAPK8 were linked to survival outcomes, suggesting a functional role for these alterations in disease progression.

PIMO’s superior performance is primarily attributable to its architectural design. PIMO incorporates interaction modeling at the earliest stages of the learning process, in contrast to DeepKEGG and PCLSurv that integrate interactions at intermediate stages. GraphPath also performs integration at early stages and consistently achieved the second-best performance, further supporting the notion that PIMO’s explicit modeling scheme enhances multi-omics analysis.

PIMO effectively captures the important genes interacting with each omics; however, explicit representation of detailed intra-omics interactions (e.g., gene-gene interactions) would be explored as a future research direction. In addition, the current pathway–pathway interaction module relies on fixed kernel sizes, which may constrain the resolution at which interactions are captured. A deeper understanding of multi-omics interactions will enable the integration of molecular signals across biological layers, providing a mechanistic foundation for more accurate patient stratification and efficient personalized therapeutic strategies in precision medicine.

## 5. Acknowledgments

This research was supported by the National Science Foundation Major Research Instrumentation (NSF MRI) (Grant#: 2117941).

